# The DC1 domain protein Vacuoleless Gametophytes regulates stamen development in Arabidopsis

**DOI:** 10.1101/2024.02.19.580957

**Authors:** Natalia L. Amigo, Leonardo A. Arias, Fernanda Marchetti, Sebastián D’Ippólito, Milagros Cascallares, Salvador Lorenzani, Jesica Frik, María Cristina Lombardo, María Cecilia Terrile, Claudia A. Casalongue, Gabriela C. Pagnussat, Diego F. Fiol

## Abstract

Vacuoleless Gametophytes (VLG) is a DC1 domain protein that was initially characterized as essential for early female and male gametophytes development in Arabidopsis. However, VLG expression was also detected in stamens, pistils and other sporophytic tissues, implying a broader role for this protein. As homozygous insertional VLG lines resulted unviable, we generated Arabidopsis amiRNA *VLG* knock-down plants to study the role of VLG in sporophyte development. The phenotypic characterization of *VLG* knock-down plants showed reduced seed set and indehiscent anthers with shorter filaments and stigma exsertion. Moreover, amiRNA *VLG* knock-down plants displayed unbroken stomia and septa, markedly reduced endothecium lignification, diminished ROS accumulation, and lower transcript levels of genes involved in jasmonic acid and lignin biosynthesis. The indehiscent phenotype was rescued by exogenous application of either jasmonic acid or H_2_O_2_. Altogether, our results suggest that VLG is involved in lignin and jasmonic acid biosynthesis pathways, and that proper levels of VLG are required in the process that leads to stomium breakage and anther dehiscence. Our findings shed light on the mechanisms underlying stamen development and provide new insights into the roles of a DC1 domain protein in plant reproduction.

**Key Message:** Vacuoleless Gametophytes participates in anther dehiscence through a process that involves JA, ROS and lignin accumulation.

## 1. Introduction

Stamen development is a complex and meticulously orchestrated process that is crucial for plant reproduction. It can be broadly divided into early stages, where cell specification occurs, and late stages, where filaments elongate, anther layers are formed (epidermis, endothecium, middle layer and tapetum) and dehiscence of anthers takes place, releasing mature pollen (Wilson et al. 2011). Each of these stages is regulated by a network of genetic and hormonal signals, and any disruption in these processes can lead to male sterility, which has significant implications for crop yield and plant biodiversity (Verma 2019).

Auxin and jasmonic acid (JA) are hormones that play pivotal roles in regulating general floral development, including stamen elongation and anther dehiscence (Cecchetti et al. 2008). Cecchetti et al. (2013, 2017) established that a peak of auxin in the middle layer negatively regulates endothecium lignification and JA biosynthesis, two fundamental processes that precede anther dehiscence. The study of numerous indehiscent mutants has shown that JA regulates the process of stomium breakage, where a specialized group of anther cell layers incur in programmed cell death allowing endothecium expansion and pollen release (Wilson et al. 2011). Moreover, JA synthesized in the upper part of the filaments is thought to regulate water transport from anthers to filaments, resulting simultaneously in anther wall dehydration and filament elongation, both crucial for a timely anthesis (Ishiguro et al. 2001).

Reactive oxygen species (ROS) also play a significant role in stamen development. During anther development, H_2_O_2_ accumulation induces lignin biosynthetic gene expression (Dai et al. 2019) and is also responsible for catalyzing lignin polymerization in the endothecium (Boerjan et al. 2003). Production of ROS may also play a role in the programmed cell death (PCD) that takes place during both septum and stomium breakage (Kurusu and Kuchitsu 2017).

Vacuoleless Gametophytes (VLG) is a DC1 domain protein required for early gametophytic development (D’ippólito et al. 2017). *VLG* mutants fail to form the central vacuole that is required early in the development of embryo sacs and pollen grains, causing early arrest in both gametophytes (D’ippólito et al. 2017). VLG is localized at the endomembrane system and it was proposed to act as a scaffold protein, as it was proved to interact with a number of proteins including SNARE associated proteins, lipases and transcription factors (D’ippólito et al. 2017). Although VLG function has only been described in gametophytes, *VLG* extensively expresses in the sporophyte, particularly in vasculature and tissues undergoing cellular expansion, which suggests that it may play additional roles in plant development (D’ippólito et al. 2017).

In this work we aimed to study the involvement of VLG during sporophyte development. We generated and analyzed *VLG* knockdown lines and we provided evidence that VLG participates in stamen development in Arabidopsis. We showed that VLG is necessary for correct anther dehiscence, in a pathway that requires JA, ROS accumulation, and lignin biosynthesis. Our findings shed light on the mechanisms underlying stamen development and provide new insights into the roles of VLG in plant reproduction.

## 2. Materials and Methods

### 2.1. Plant materials and growth conditions

*Arabidopsis thaliana* ecotype Columbia (Col-0) WT and knock-down plants were grown in growth chambers under long-day conditions (16-h light/8-h dark) at 25°C and 50-70% humidity. Seeds were treated in 10% (v/v) sodium hypochlorite, washed five times with sterile water, stratified at 4°C for 48 h and plated on Murashige and Skoog media, with 50 μg/ml kanamycin and/or 50 µg/ml hygromycin when necessary, at the light and temperature conditions indicated above. Resistant seedlings were subsequently moved from plates to soil. Arabidopsis plants were transformed using the floral dip method (Clough and Bent 1998) with the *Agrobacterium tumefaciens* strain GV3101. proVLG:GUS plants were previously reported by (D’ippólito et al. 2017), The numerical designation of developing flower buds was adopted according to (Smyth et al. 1990).

### 2.2. Knock-down of VLG

VLG knock-down plants were generated using an artificial microRNA (amiR) strategy described in (Schwab et al. 2010). Primers listed in Table S1 were designed using Web MicroRNA Designer (WMD3, http://wmd3.weigelworld.org/cgi-bin/webapp.cgi) tools to incorporate a 21-bp amiRNA sequence into the MIR319a vector. Subsequently, the amiRNA construct was subcloned into the expression vector pCHF3. The resulting amiR-VLG construct was transformed into Arabidopsis as indicated in the previous section and independent lines were selected (VLG-amiRNA L1-L3).

### 2.3. Morphological and histological analyses

Flowers were collected before anthesis and dissected and cleared in Hoyer’s solution or 90% ethanol overnight. After clearing, inflorescences and anthers were observed on a Zeiss Axioplan Imager-A2 microscope under differential interference contrast (DIC) optics. Images were captured on an Axiocam HRC CCD camera (Zeiss) using the Axiovision program (version 4.2). For GUS staining, collected proVLG:GUS developing seedlings and inflorescences were processed as described by (Pagnussat et al. 2007).

### 2.4. Light and confocal microscopy and lignin detection

Inflorescences were fixed in a solution of ethanol:water:phormol:acetic acid (10:7:2:1) and then dehydrated and included in Paraplast (Sigma) according to (D’Ambrogio de Argüeso, 1986, Ruzin 1999). Samples were mounted and cut in a rotary microtome. The sections were stained with Toluidin Blue O and were examined by bright field microscopy in a Nikon Eclipse E 2000 microscope (Tokyo). Images were obtained with a digital camera (Nikon Coolpix 990).

Confocal microscopy was performed using a BX61-FV300 microscope (Olympus Optical, Tokyo, Japan) with an acridine orange/ethidium bromide stain, according to (Yang et al. 2007). The acridine orange and ethidium bromide fluorescence intensities (green and red respectively) were digitally quantified from 30 anthers of each condition using Image J software (Fiji v.2.15.0).

### 2.5. In situ determination of reactive oxygen species

Detection of hydrogen peroxide (H_2_O_2_) in plant tissues was performed by DAB staining according to (Villarreal et al. 2009). Briefly, inflorescences of Arabidopsis plants were incubated overnight in total darkness with a DAB solution (0.5 mg/ml) of 3,30-diaminobenzidine (Sigma) in 50 mM sodium acetate buffer (pH 5, 4) at 25°C. Afterward, the solution was removed and samples were decolored with 90% ethanol overnight at 25°C to remove chlorophyll. A dark brown color was visualized for H_2_O_2_ due to the oxidation of DAB. Images were taken with a binocular SMZ800 (Nikon) and a CoolSnap Pro camera (MediaCybernetics, TX). The DAB signal was digitally quantified as the average pixel light value of about 40 anthers from each condition using Image J software (Fiji v.2.15.0).

### 2.6. RNA isolation and RT qPCR analysis

Total RNA was isolated either from inflorescences (pooled flower buds from stages 1-14) or from anthers (about 120 anthers pooled from stages 1-11) with TRIzol reagent (Invitrogen, USA). The amount and quality of RNA were checked by spectrophotometry (OD: 260/280) and by electrophoresis in agarose gels. cDNA was obtained from 1 μg of total RNA pretreated with DNase I (Promega) by reverse transcription reaction using ImProm-II Reverse Transcriptase (Promega) with random hexamers primers. qPCR reactions were performed using FastStart Universal SYBR Green Master (ROX) PCR mix (Roche) in a StepOne machine (Applied Biosystems). The amplification conditions were 95°C for 10 min, followed by 40 cycles of amplification (95°C for 15 s, 60°C for 30 s) and reactions were conducted in triplicates. Data presented are normalized to the expression level of the control gene actin2 of three independent experiments. Sequences of the primers used are listed in Table S2. The ‘delta–delta Ct method’ was used to calculate the relative expression of the genes (Livak and Schmittgen 2001).

### 2.7. Application of methyl jasmonate (MeJA) and H_2_O_2_

All opened flowers (after stage 14) were removed from the inflorescence, and the remaining flower bud clusters were dipped into 1mM MeJA (Sigma-Aldrich) or 1mM H_2_O_2_ dissolved in 0.05% aqueous Tween-20 as indicated in (Shih et al. 2014). The plants were then grown in a growth room under a long day (16/8 h, light/dark) photoperiod at 25°C for 40-50 days. The development of the silique for MeJA or H_2_O_2_ treated flower buds was recorded and pictures were taken 3 days and 30 days after treatment.

### 2.8. Statistical analysis

Independent experiments were conducted at least three times and assays were performed at least in triplicate. The values obtained were used for statistical analysis between groups using GraphPad Prism, version 6.01. Statistical differences between groups were calculated by applying the Student’s t-test or analysis of variance test; and Tukey’s or Dunnett’s multiple comparisons test were performed, considering P < 0.05 as significant.

## 3. Results

### 3.1. VLG knock-down causes reduced seed set, anther indehiscence and impaired anthesis

VLG is essential for early gametophyte development and the *vlg* mutant phenotype is highly penetrant, making homozygous lines unviable (D’ippólito et al. 2017). Therefore, to assess VLG function in the sporophyte, we generated a construction expressing *pro35S:amiRNA* to specifically target *VLG* transcripts and obtained three knock-down transgenic lines (amiR-VLG L1-3). We confirmed *VLG* knocked-down expression in inflorescences from L1, L2 and L3 lines; VLG transcripts were reduced by 68%, 83% and 84%, respectively (Fig. 1A).

**Fig 1.**
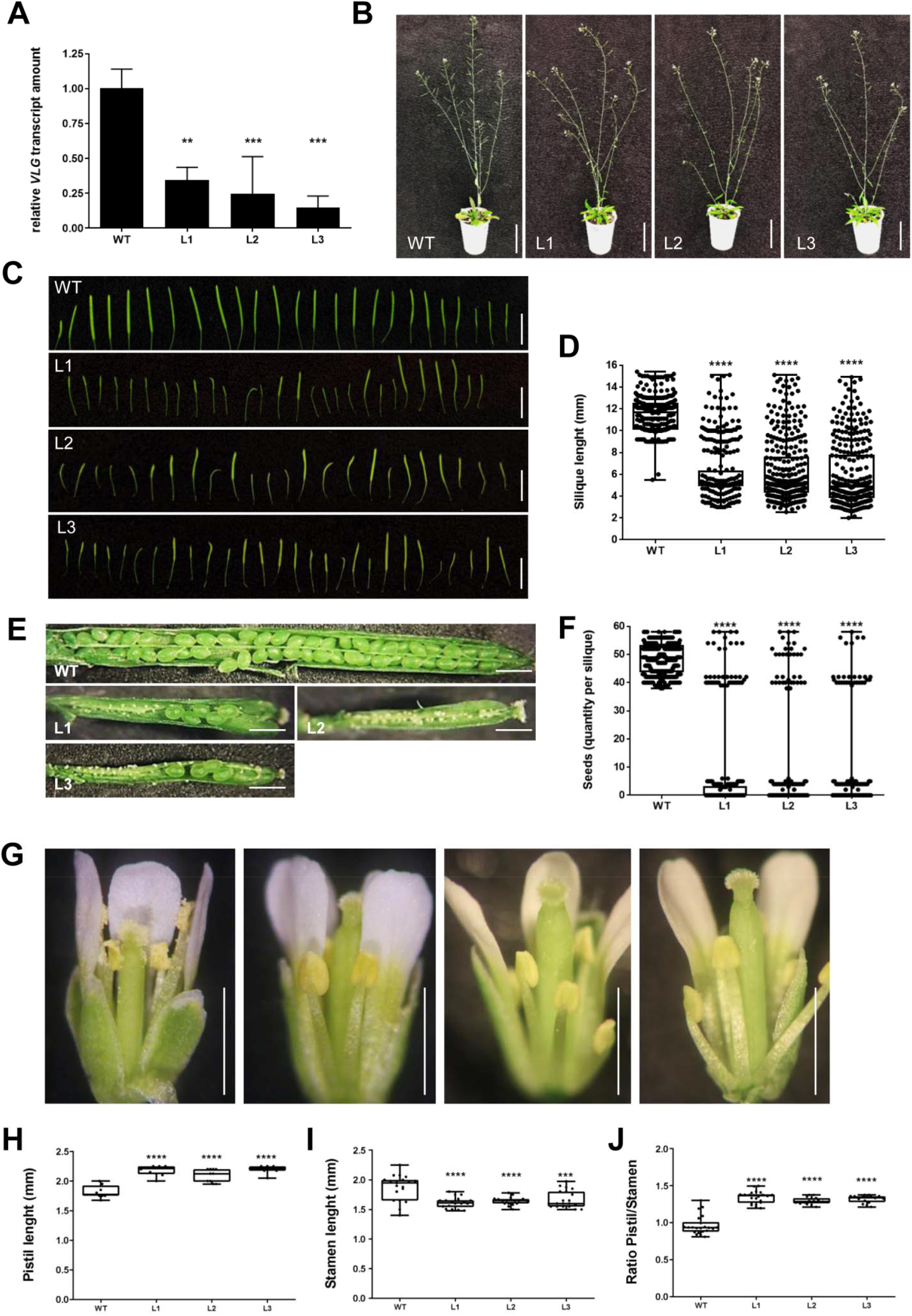
Analysis of amiR-VLG plants. Relative quantification of VLG transcripts in anthers (stages 1 to 11) according to (Smyth et al., 1990) from WT plants and amiR-VLG lines L1, L2 and L3 **(A)**. 42-day-old WT and amiR-VLG (lines L1, L2 and L3) plants **(B)**. Bars: 50 mm. Collection of main stem siliques displayed in chronological order, from the earliest ones to the latest ones **(C)** and silique length quantification **(D)** of WT plants and amiR-VLG lines L1, L2 and L3. (n ≥ 320 siliques from each line). Bars: 10 mm. Seed sets **(E)** and quantification of seeds per silique **(F)** of WT plants and amiR-VLG lines L1, L2 and L3. (n ≥ 275 siliques from each line). Flowers at stage 15 **(G)** and their corresponding pistil length **(H)**, stamen length **(I)**, and ratio of pistil/stamen **(J)** from WT plants and amiR-VLG lines L1, L2 and L3. (n ≥ 20 flowers from each line). Bars: 10mm. Asterisks indicate significant differences compared to WT plants. ANOVA and Dunnett’s multiple comparisons test, p<0.001 (***), p < 0.0001 (****).

amiR-VLG plants developed normally until flowering, except for minor differences in rosette leaves, which displayed an epinastic phenotype (curved towards their abaxial side) under normal growth conditions (Fig. 1B, Fig S1A). At flowering, all three amiR-VLG lines consistently presented shorter siliques, with mean values of 6.06 mm, 7.02 mm and 6.57 mm for L1, L2 and L3 respectively, compared to a mean of 10.82 mm for WT plants (Fig. 1C and D). Concomitantly, a lower number of seeds was present in the siliques, with mean values ranging from 5 to 9 in amiR-VLG lines, compared to about 48 for WT siliques (Fig, 1E and F). This phenotype was more marked in the first 8-10 siliques and became less pronounced as the plants aged, as it can be seen in Fig. 1C.

To better assess the causes of the reduced seed set, we performed a morphological characterization of floral structures. Flowers from amiR-VLG plants were similar to WT flowers regarding opening time, sepals, petals and development of stigmatic papillae. However, amiR-VLG flowers presented indehiscent anthers, significantly shorter stamen filaments (around 12% shorter than filaments from WT plants) and longer pistils (about 19% longer than pistils from WT plants) (Fig. 1G to I) thus, resulting in a substantially higher pistil/stamen length ratio (Fig. 1J) that likely hinders pollination.

To inspect the cause of the amiR-VLG-related indehiscent phenotype, we analyzed transverse sections of anthers during anthesis (stage 14). As opposed to WT anthers, amiR-VLG anthers did not show septa and stomia breakage (Fig. S1C). Additionally, indehiscent anthers also appeared collapsed, suggesting a weak secondary wall structure (Fig. S1C).

### 3.2. VLG is expressed throughout flower development

In order to assess whether VLG could be involved in flower development, we analyzed the expression profile of *proVLG:GUS* reporter lines. VLG promoter activity is detectable all along the flower development process (Fig. 2). During the early stages of flower buds, from stages 1 to 10 (according to Smyth et al., 1990), the expression occurs in all flower organs (Fig. 2B to F). However, as the flower matures, it becomes more localized to the anthers, stamen filaments, and style (Fig. 2G to P).

**Fig 2.**
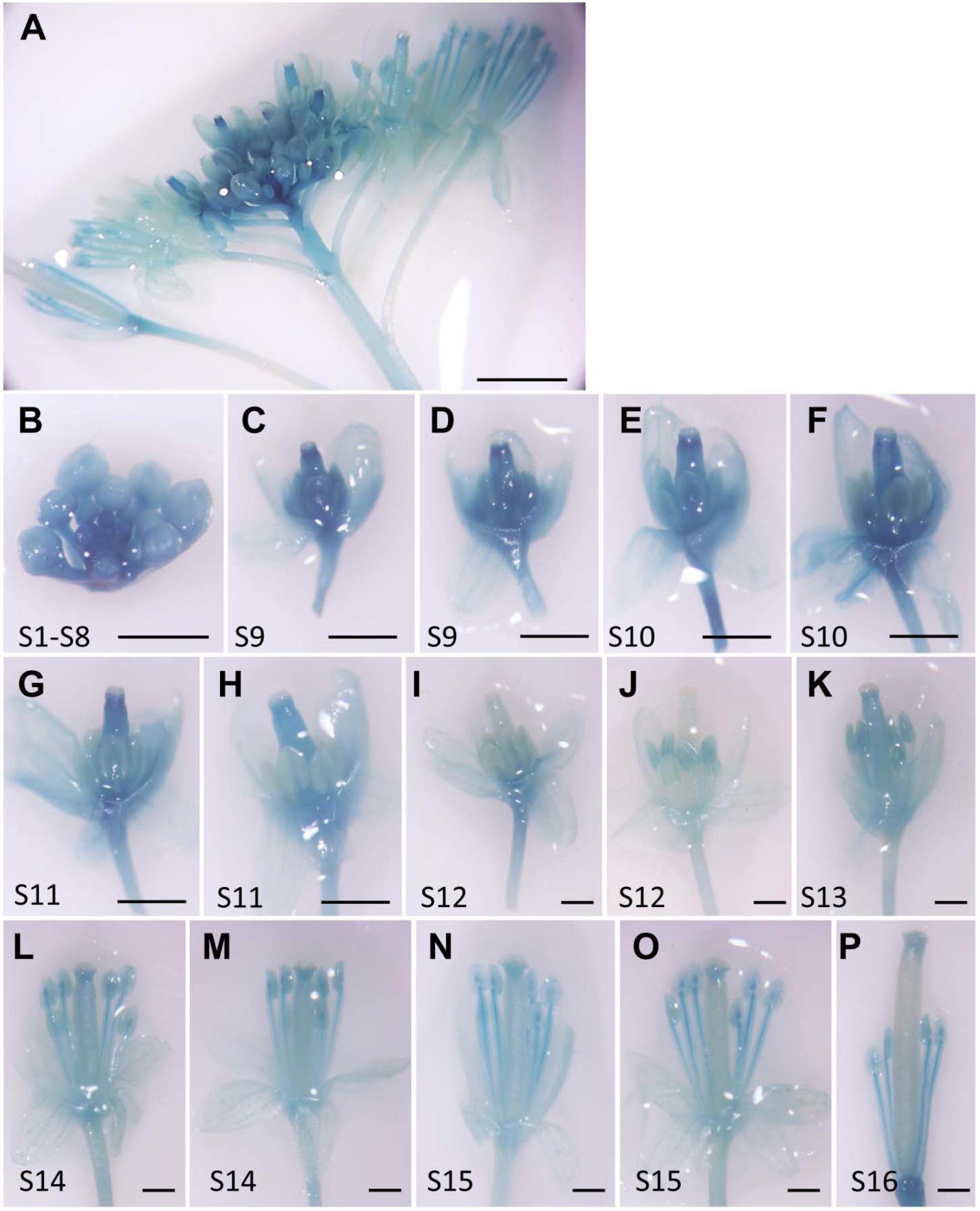
*VLG* promoter activity in inflorescences. **(A)** GUS signal was detected in inflorescences from 5-week-old proVLG:GUS plants. Different stages of inflorescence are indicated: **(B)** stages 1 to 8, **(C-D)** stage 9, **(E-F)** stage 10, **(G-H)** stage 11, **(I-J)**, stage 12, **(K)** stage 13, **(L-M)** stage 14, **(N-O)** stage 15, **(P)** stage 16. Three independent transgenic lines were used for this study. Pictures are representative of the results obtained in all analyzed lines. Bars: A, 5 mm; B-K, 0.5 mm; L-P, 1 mm.

### 3.3. VLG knock-down lines show impaired lignification and reduced H_2_O_2_ accumulation in anthers

Several anther indehiscent phenotypes have been reported in the literature, either caused by impaired endothecium lignification (Cecchetti et al. 2013) or reduced H_2_O_2_ accumulation (Dai et al. 2019). To analyze endothecium lignification, we used confocal microscopy to examine anthers stained with acridine orange/ethidium bromide to highlight primary cell walls and lignification respectively. We observed that primary cell wall content, as detected by acridine orange staining, did not differ in intensity between WT and amiR-VLG lines; however, it evidenced that the shape of endothecium cells in amiR-VLG lines was heavily distorted (Fig. 3A and B). WT cells are roughly even in size and present an orthogonal shape with a wide contact surface between cell walls. Conversely, amiR-VLG endothecium cells are highly irregular in size, rounder, and display narrow contact surfaces (Fig. 3A). On the other hand, it was observed a very significant reduction of lignin content in anthers from amiR-VLG lines L1, L2 and L3 by acridine orange staining (Fig. 3A and C).

**Fig 3.**
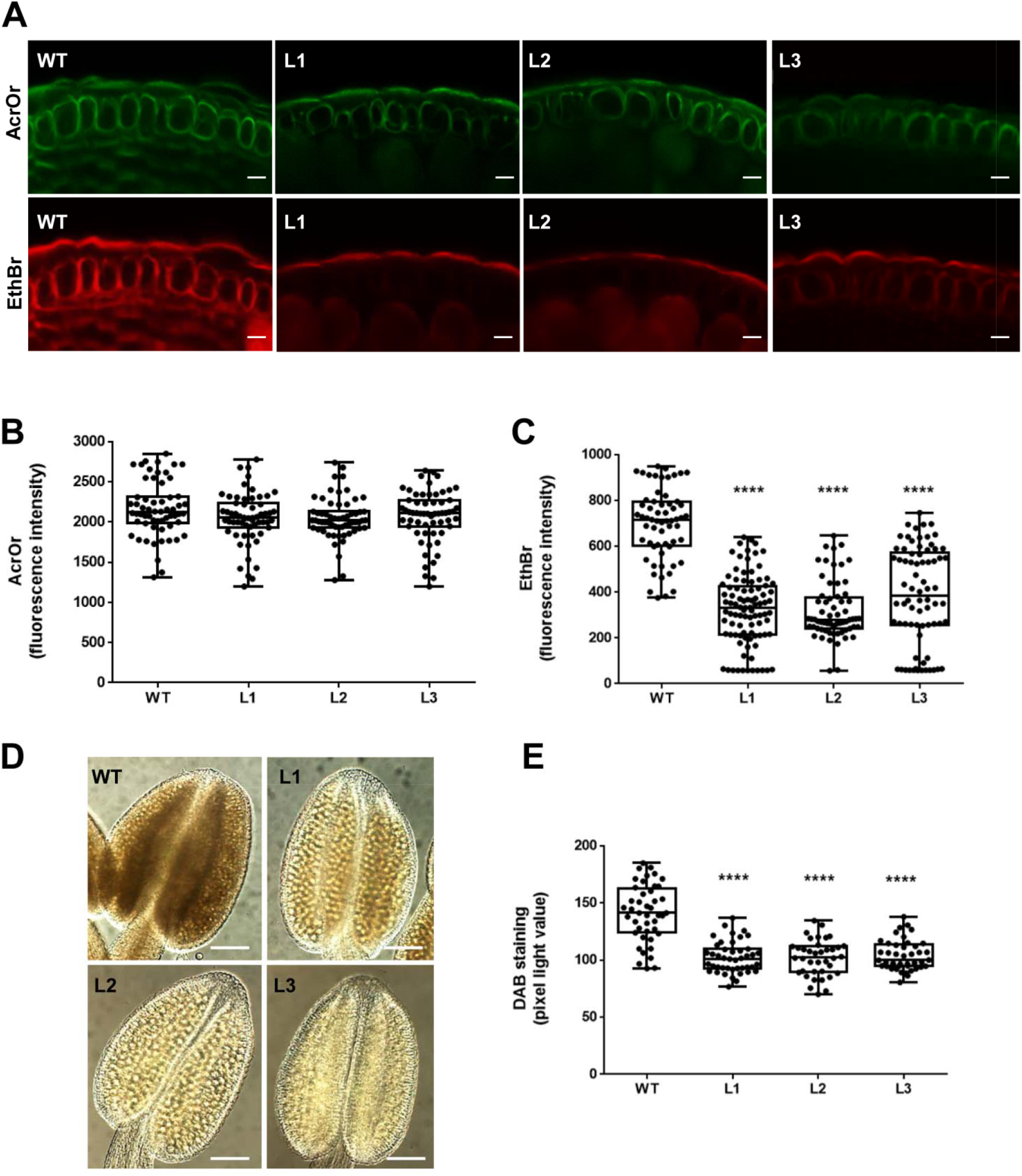
Anther development is affected in amiR-VLG lines. Anthers at stages 11 to 13 from WT plants and amiR-VLG lines (L1, L2 and L3) were stained with acridine orange/ethidium bromide to show the distribution of primary cell walls (AcrOr, green fluorescence, upper panels) and lignin (EthBr, red fluorescence, lower panels) **(A)**. Quantification of green **(B)** and red **(C)** fluorescences representing primary cell walls and lignified walls, respectively. Bars: 10 µm. Anthers at stages 11 to 13 from WT plants and amiR-VLG lines (L1, L2 and L3) were analyzed by DAB staining to detect **(D)** and quantify **(E)** ROS accumulation. Bars: 0.1 mm. ANOVA and Tukey’s multiple comparisons test, p < 0.0001 (****).

To assess if H_2_O_2_ accumulation is involved in the reduced lignification observed, we used diaminobenzidine (DAB) staining on whole anthers from WT and amiR-VLG plants. We examined anthers between stages 11 and 13, as these stages were reported to be relevant for H_2_O_2_ accumulation by Dai et al. (2019). Interestingly, we found a marked decrease in H_2_O_2_ accumulation in all three amiR-VLG lines, showing DAB staining values about one-third lower compared to WT anthers (Fig. 3D and E).

### 3.4. Knock-down of VLG reduces the expression levels of JA and lignin biosynthetic genes

As amiR-VLG lines displayed phenotypes consistent with impaired lignin and JA biosynthesis, we analyzed the expression of key genes belonging to those pathways in anthers (pooled from stages 1 to 11) from WT and amiR-VLG lines. Thus, we measured transcript abundance through RT qPCR of *SND2* (*Secondary wall-associated NAC Domain 2, NAC073*), a gene that encodes a transcription factor that regulates the expression of genes involved in secondary cell wall development (Zhong et al. 2008) and three cellulose biosynthesis genes: *CESA4* (*Cellulose synthase A 4*) (Richmond and Somerville 2000), *IRX3* (*Irregular Xylem 3*) and *IRX12* (*Irregular Xylem 12*) (Hussey et al. 2011). We found significantly lower transcript levels in these four genes in the three amiR-VLG lines compared to WT plants (Fig. 4A), supporting the hypothesis that VLG is involved in the regulation of lignin biosynthesis.

**Fig 4.**
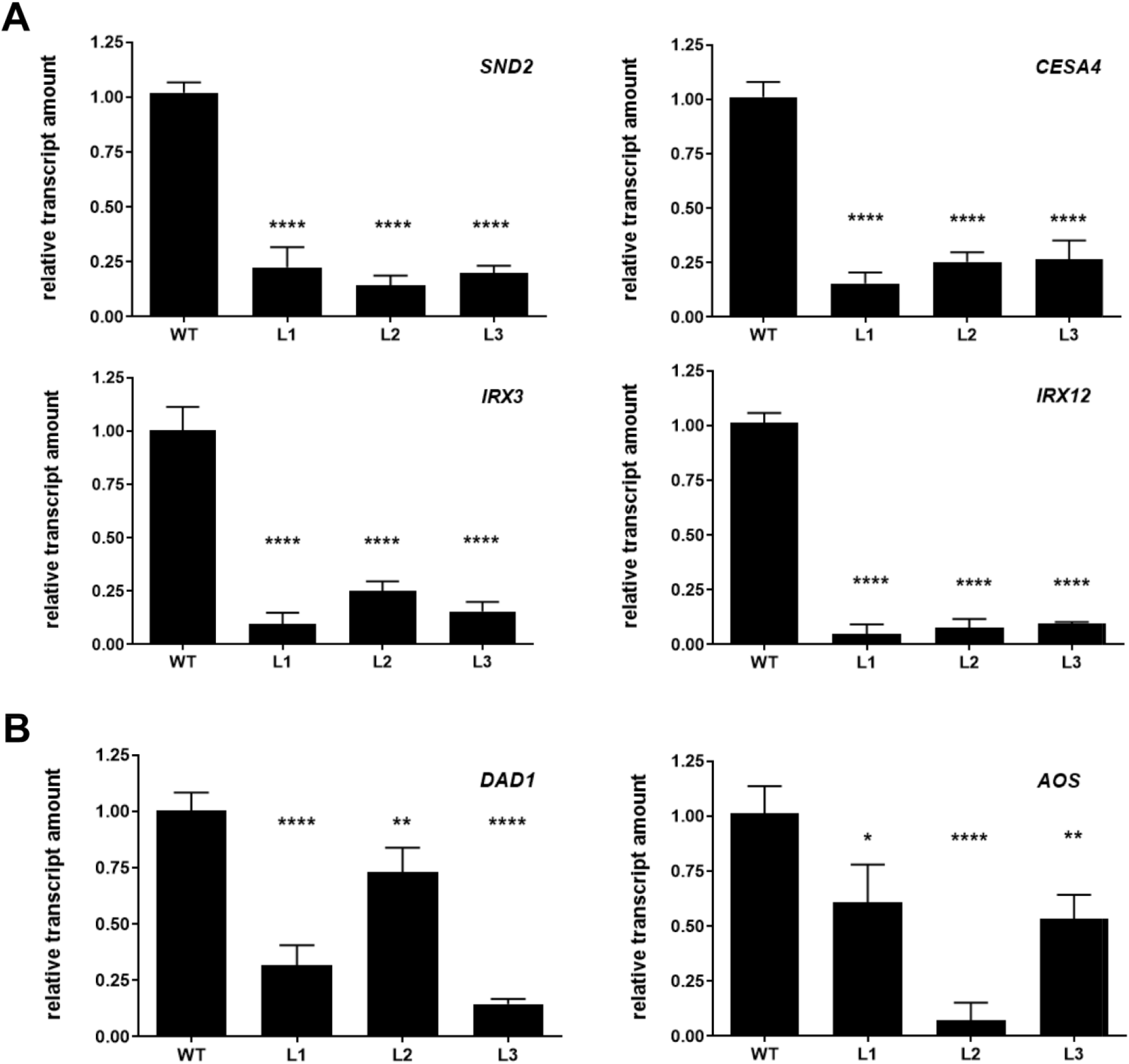
Transcript levels from lignin and JA biosynthesis pathways are affected in amiR-VLG lines. Relative transcript levels were determined by RT qPCR from total RNA samples isolated from anthers (pooled from stages 1 to 11) of 5-week-old WT and amiR-VLG lines L1, L2 and L3. Genes from lignin biosynthesis pathway: *SND2 (Secondary wall-associated NAC Domain 2*), *CESA4* (*Cellulose synthase A4*), *IRX3 (Irregular Xylem 3)* and *IRX12 (Irregular Xylem 12)* **(A)**. Genes from JA biosynthesis pathway: *DAD1* (*Defective in Anther Dehiscence 1)* and *AOS* (*Allene Oxide Synthase)* **(B)**. Data are means of three replicates ± SD. Asterisks indicate significant differences in comparison to WT plants by ANOVA and Dunnett’s multiple comparisons test, p<0.05 (*), p<0.01 (**), p<0.001 (***), p < 0.0001 (****).

In addition, we determined the abundance of two gene transcripts involved in JA biosynthesis, *DAD1* (*Defective in Anther Dehiscence 1*) which encodes a lipolytic enzyme that catalyzes the synthesis of linolenic acid in the initial step in the JA biosynthetic pathway (Tabata et al. 2010) and *AOS* (*Allene Oxide Synthase, CYP74A*) which encodes a cytochrome P450 enzyme that catalyzes the dehydration of hydroperoxide to an unstable allene oxide in the same pathway (Park et al. 2002). We also found significantly lower transcript levels of these two genes in the three amiR-VLG lines compared to WT plants (Fig. 4B), suggesting that *VLG* could be involved in the regulation of the JA biosynthetic pathway.

### 3.5. amiR-VLG anther indehiscent phenotype is rescued by MeJA and H_2_O_2_ exogenous treatments

In previous studies, authors have been able to rescue indehiscent anthers by exogenous applications of jasmonates (Wilson et al. 2011) or H_2_O_2_ (Dai et al. 2019). To test if amiR-VLG indehiscent lines were defective in a pathway dependent on JA or H_2_O_2_, we applied MeJA (1 mM) or H_2_O_2_ (1 mM) by briefly immersing inflorescences in either solution once a day for three consecutive days. Strikingly, each treatment independently restored correct anther dehiscence in all three amiR-VLG lines, resulting in proper fruit development with a complete seed set (Fig. 5 and Fig. 6). The mean quantity of seeds per silique upon MeJA treatment changed from 7.3 to 45.4 (amiR-VLG L1), from 6.3 to 46.6 (amiR-VLG L2) and from 6.1 to 45.4 (amiR-VLG L3), compared to WT, which remained in the same range, changing from 48.4 to 48.0 (Fig. 5). The mean quantity of seeds per silique upon H_2_O_2_ treatment changed from 5.3 to 43.4 (amiR-VLG L1), from 9.3 to 44.0 (amiR-VLG L2) and from 4.4 to 45.6 (amiR-VLG L3), compared to WT, which remained about 48.5 in both conditions (Fig. 6). These results suggest that VLG is required to maintain proper levels of JA and H_2_O_2_ in anthers. In contrast, none of these treatments reverted either the short filament or the long pistil phenotypes of amiR-VLG lines, even when anther dehiscence was indistinguishable from WT plants (Fig. 5A and B, Fig. 6A and B and Fig. S2).

**Fig 5.**
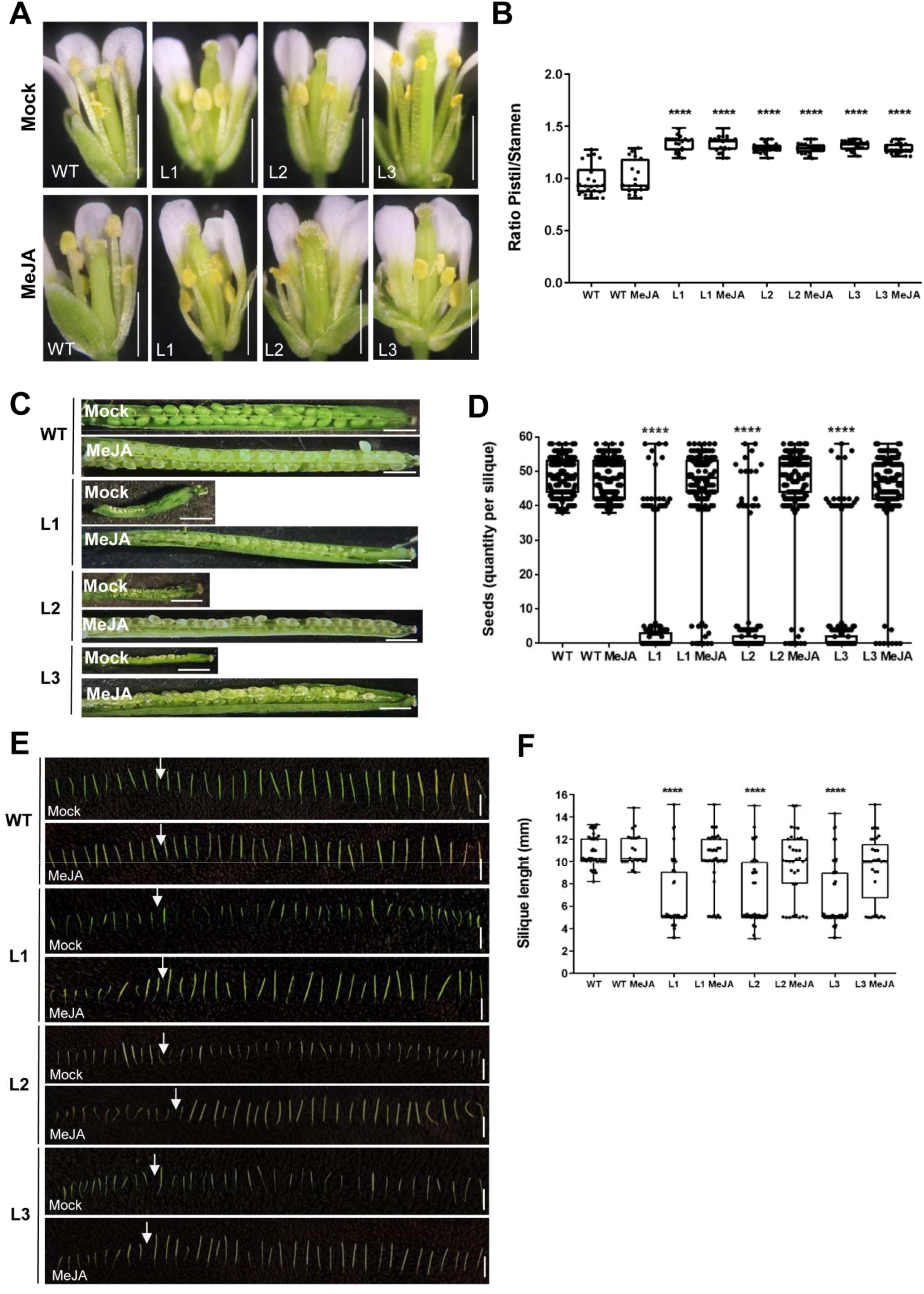
Indehiscence of anthers is rescued by Methyl Jasmonate (MeJA) exogenous application in amiR-VLG plants. Flowers in 4-week-old WT and amiR-VLG (lines L1, L2 and L3) plants were treated with 1 mM MeJA for three days. Three days after the last application flowers were pictured **(A)** and ratio of pistil/stamen was measured (n ≥ 20 flowers from each line) **(B)**. Thirty days after treatment, seed content was pictured **(C)** and quantified (n ≥ 175 siliques from each line) **(D)** and silique length was pictured **(E)** and quantified (n ≥ 150 siliques from each line) **(F)**. Arrows indicate the treatment starting point. Bars: 10 mm. Asterisks indicate significant differences compared to WT control plants. ANOVA and Dunnett’s multiple comparisons test, p < 0.0001 (****).

**Fig 6.**
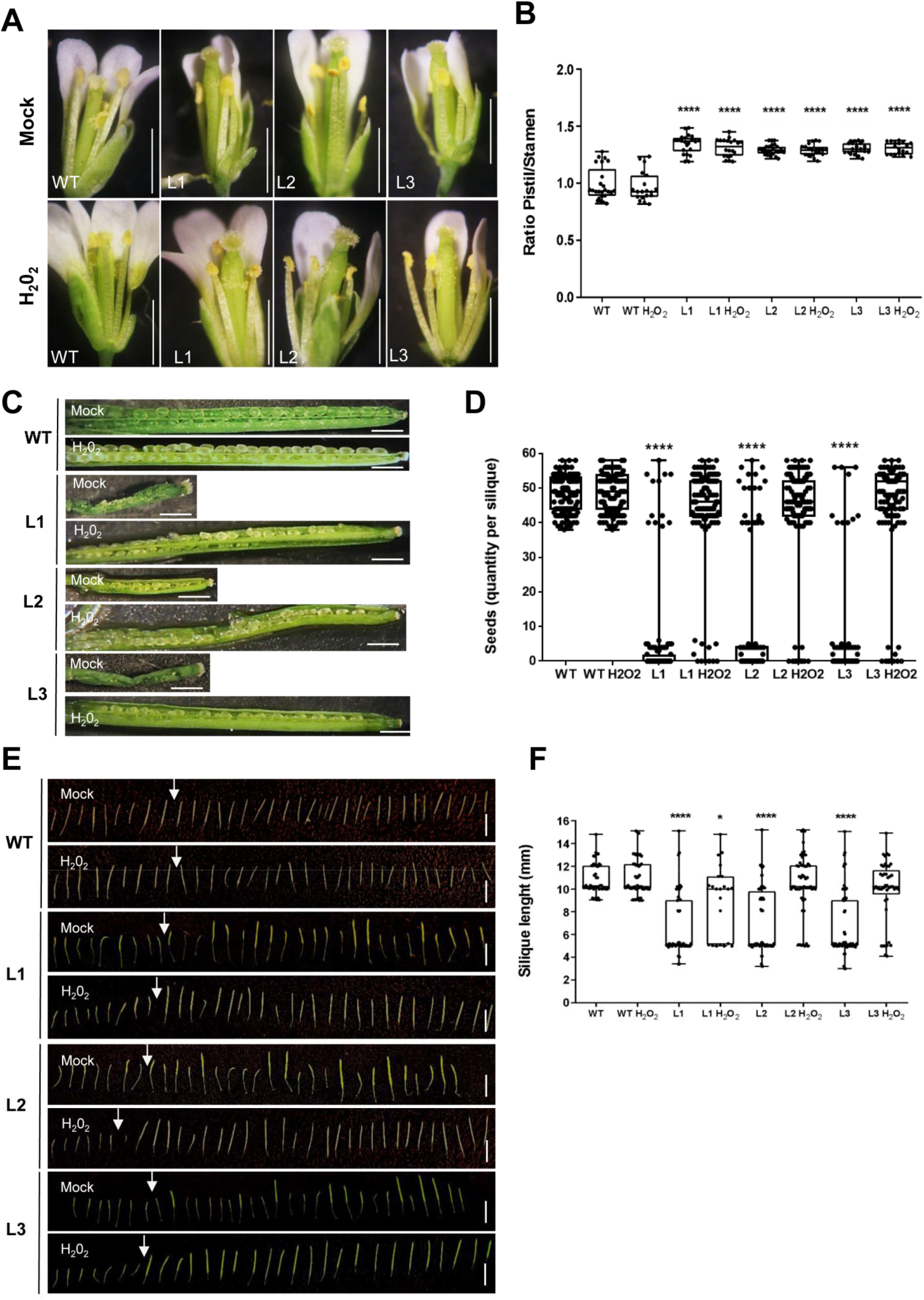
Indehiscence of anthers is rescued by H_2_O_2_ exogenous application in amiR-VLG plants. Flowers in 4-week-old plants WT and amiR-VLG (lines L1, L2 and L3) plants were treated with 1 mM H_2_O_2_ for three days. Three days after the last application flowers were pictured **(A)** and ratio of pistil/stamen was measured (n ≥ 20 flowers from each line) **(B)**. Thirty days after treatment, seed content was pictured **(C)** and quantified (n ≥ 120 siliques from each line) **(D)** and silique length was pictured **(E)** and quantified (n ≥ 50 siliques from each line) **(F)**. Arrows indicate the treatment starting point. Bars: 10 mm. Asterisks indicate significant differences compared to WT control plants. ANOVA and Dunnett’s multiple comparisons test, p<0.05 (*), p < 0.0001 (****).

We next investigated the effect of MeJA and H_2_O_2_ exogenous treatments on the expression of the sets of genes related to the secondary wall and JA biosynthetic pathways. Interestingly, we observed that MeJA treatment was able to restore the expression levels of the genes of both pathways in the anthers (pooled from stages 1 to 11) in all three amiR-VLG lines (Fig. S3). On the other hand, H_2_O_2_ treatment did not affect the expression levels of either the secondary wall or the JA biosynthetic pathway genes in the amiR-VLG lines (Fig. S4).

## 4. Discussion

In this work we studied the role of VLG in the sporophytic development of Arabidopsis and found that it is required for the proper development of the stamen. Knocking down *VLG* via amiRNA led to anther indehiscence, short filaments and stigma exsertion, causing a drastic decrease in the seed set (Fig. 1). Using lines carrying the *proVLG:GUS* reporter, we showed that *VLG* is expressed throughout all the developmental stages of the flower, exhibiting strong expression levels in filaments and during early stages of anther development (Fig. 2). This expression pattern, coupled with the male sterility phenotype observed in amiR-VLG lines, suggests that VLG may play a role in the processes of filament elongation and anther dehiscence.

For successful fertilization, flowers must undergo a series of developmental events in a timely manner. These events include the development of the pistil and stigmatic papillae, the growth of stamens, and the opening of anthers to release pollen. These events are regulated by a combination of genetic, hormonal, and environmental factors. The accurate and timely dehiscence of anthers relies on several key steps, including the development of the four anther cell layers, endothecium lignification, tapetum degeneration, and the opening of septa and stomia (Wilson et al. 2011). The regulation of endothecium lignification is governed by both hormonal control and ROS balance. At floral stage 10, an auxin peak negatively regulates endothecium lignification via MYB26, a master activator of cell wall secondary thickening (Cecchetti et al. 2013). In fact, auxin suppresses NAC family transcription factors SND1/2, which directly target and induce MYB26 (Xiao et al. 2021). Auxin also negatively regulates JA biosynthesis by inducing ARF6/8 transcription factors, which indirectly inhibit DAD1, the initial enzyme in the JA biosynthetic pathway (Tabata et al., 2010). After reaching its peak, auxin concentration decreases, and the repression of MYB26 is lifted, enabling MYB26 to initiate endothecium lignification. This reduction in auxin also promotes higher expression levels of DAD1 and OPR3, resulting in an increase in JA concentration. Consequently, this leads to stomium breakage and the completion of the anther dehiscence process at stage 13.

We performed transverse sections of anthers before anthesis, revealing the absence of septum and stomium rupture and an apparent lack of endothecium lignification, and confirmed also decreases in lignin contents in the amiR-VLG plants (Fig. S1B, Fig. 3A and C).

The breakage of the stomium is triggered by JA. In most JA biosynthesis mutants, the absence of stomium breakage phenotype can be reversed by MeJA application (Wilson et al., 2011). To ascertain whether the JA biosynthesis pathway was affected in amiR-VLG lines, we quantified two transcripts of the JA biosynthetic pathway, *DAD1* and *AOS*, and observed a significant reduction in the abundance of both transcripts in anthers (pooled from stages 1 to 11) of amiR-VLG lines L1, L2 and L3 (Fig. 4B). In agreement with a possible role for VLG regulating JA levels in anthers, we successfully rescued the indehiscent phenotype of amiR-VLG plants with applications of MeJA (Fig. 5). Thus, we infer that the phenotype of indehiscent anthers observed in amiR-VLG lines is consistent with a role for VLG in JA biosynthesis; VLG is required to achieve JA levels necessary to trigger anther opening. In agreement with that, previous studies have shown that reduced expression of AOS and DAD1 can lead to male sterility by blocking JA synthesis (Park et al. 2002, Sanders et al. 2000). We also found that MeJa treatment was able to increase AOS and DAD1 transcript levels in amiR-VLG lines L1, L2 and L3 (Fig. S3B), complementing the reduction caused by VLG knock down.

Endothecium secondary thickening, and in particular lignin deposition which occurs at stage 11, are essential for proper anther dehiscence (Cecchetti et al, 2017, Wilson et al, 2011). The final polymerization steps of lignin biosynthesis involve the activation of monolignols to free radicals, which are mediated by oxidation systems such as peroxidase/H_2_O_2_, followed by non-enzymatic coupling of monolignol radicals to make the final lignin polymers (Boerjan et al. 2003). Thus, proper levels of H_2_O_2_ are required for lignin accumulation in the endothecium (Dai et al. 2019). We observed that amiR-VLG plants presented reduced ROS accumulation in anthers (Fig. 3D and E) and exogenous application of H_2_O_2_ on floral buds successfully restored proper stomium breakage (Fig. 6), suggesting that VLG could be involved in lignin accumulation by regulating H_2_O_2_ levels. Anthers from amiR-VLG lines also showed a decrease in transcript levels of genes involved in secondary cell wall biosynthesis (*SND2, CESA4, IRX3* and *IRX12*) (Fig. 4A); however, the application of H_2_O_2_ did not affect the expression of these genes (Fig. S4). We hypothesize that the addition of H_2_O_2_ was able to compensate for the reduced ROS levels in amiR-VLG lines, leading to higher lignin polymerization and increased lignin deposition in treated anthers, restoring this way the dehiscence phenotype, despite the lower expression of the lignin biosynthetic pathway genes.

Remarkably, SND2 was previously reported as a VLG interacting protein, as detected by a yeast two-hybrid assay (D’ippólito et al. 2017), suggesting that VLG could also participate in the lignin biosynthetic process by interacting with the master regulator gene of secondary thickening. Interestingly, the interaction between another DC1 protein (binucleate pollen, BNP) and two other NAC family transcription factors (Vascular Plant One-Zinc Finger 1 and 2, VOZ1/2) was recently reported (Arias et al. 2022). In this case, it was proposed that BNP, which is localized as VLG in the endomembrane system, was required for a timely nuclear translocation of VOZ1/2 during pollen development (Arias et al. 2022).

VLG knocked-down plants treated with either MeJA or H_2_O_2_ did not recover the short filament phenotype (Fig. 5 and Fig. 6). Filament elongation is hypothesized to rely upon JA synthesized in the upper regions of the filaments, which regulates water transport from the anthers to the filaments (Ishiguro et al. 2001). However, this process appears to be uncoupled from anther dehiscence (Cecchetti et al. 2008). Therefore, the timing and concentrations of exogenous applications required to rescue those phenotypes may differ substantially. During the late phase of stamen development, other molecules, such as auxins, play a crucial role in controlling the process. For instance, the maximum concentration of auxin is established in the anthers at stage 10; this triggers filament elongation and negatively controls anther dehiscence and pollen maturation. As tapetum degenerates, the concentration of auxin decreases. Misregulation in the occurrence or in the timing of the components involved in the anther development process can lead to a reduced seed set in amiR-VLG lines. To better understand the involvement of this important hormone in the process, a study of auxin distribution in different stages of stamen development in VLG mutants will be necessary.

Based on these findings and existing literature, we propose that VLG modulates the expression of JA biosynthesis genes, leading to timely stomium breakage. VLG is also involved in upstream H_2_O_2_ accumulation in anthers around flower stage 11. The peak of H_2_O_2_ in turn induces lignin polymerization, resulting in proper endothecium lignification. A possible way of action of VLG in the lignin biosynthesis pathway could be through the interaction with SND2, the secondary wall thickening master regulator. Deficient VLG levels in amiR-VLG stamens could impair the formation of molecular complexes required for JA biosynthesis, H_2_O_2_ accumulation and lignin biosynthesis required for proper anthesis. Follow-up work will be aimed to identify the proteins interacting with VLG in this process.

## Acknowledgments

We thank Viviana Daniel for her technical assistance with confocal microscopy.

