## Supplemental Figs..S1-4, Tables S1-2 for "The DC1 domain protein Vacuoleless Gametophytes regulates stamen development in Arabidopsis"

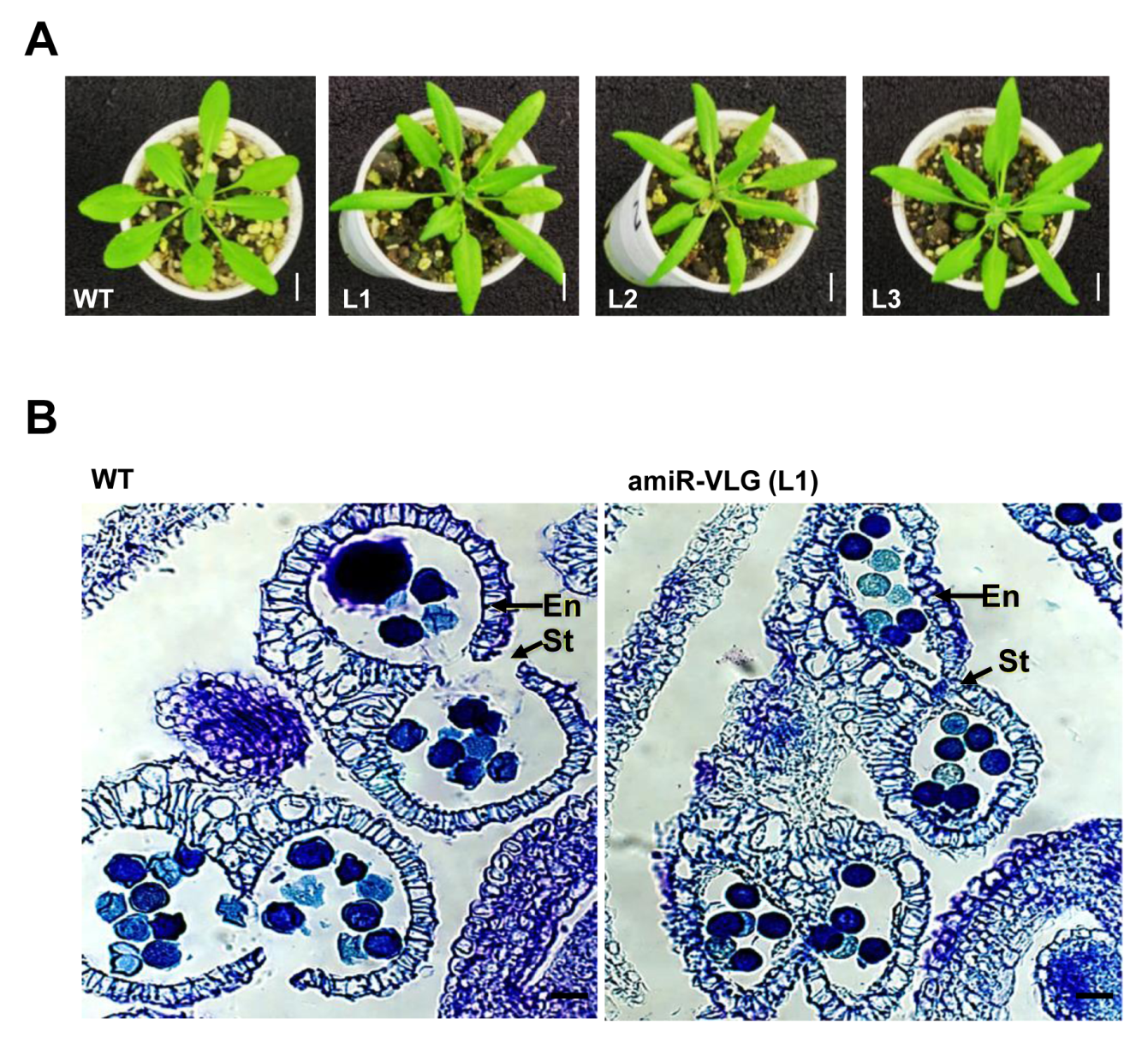

Fig. S1

**Fig. S1.** **Analysis of amiR-VLG lines**

26-day-old WT and amiR-VLG lines L1, L2 and L3 rosettes showing leaf epinasty phenotypes **(A)**. Bars: 50mm.

amiR-VLG anthers show unbroken stomium **(B)**. Micrographs of transverse sections of anthers from WT plants (displaying normal structure and stomium breakage) and amiR-VLG (L1) plants (displaying unbroken stomium and colapsed locules). Arrows indicate the site for stomium breakage (open in WT, not open in amiR-VLG L1). St: stomium. En: endothecium. Bars: 20 µm.

Fig. S2

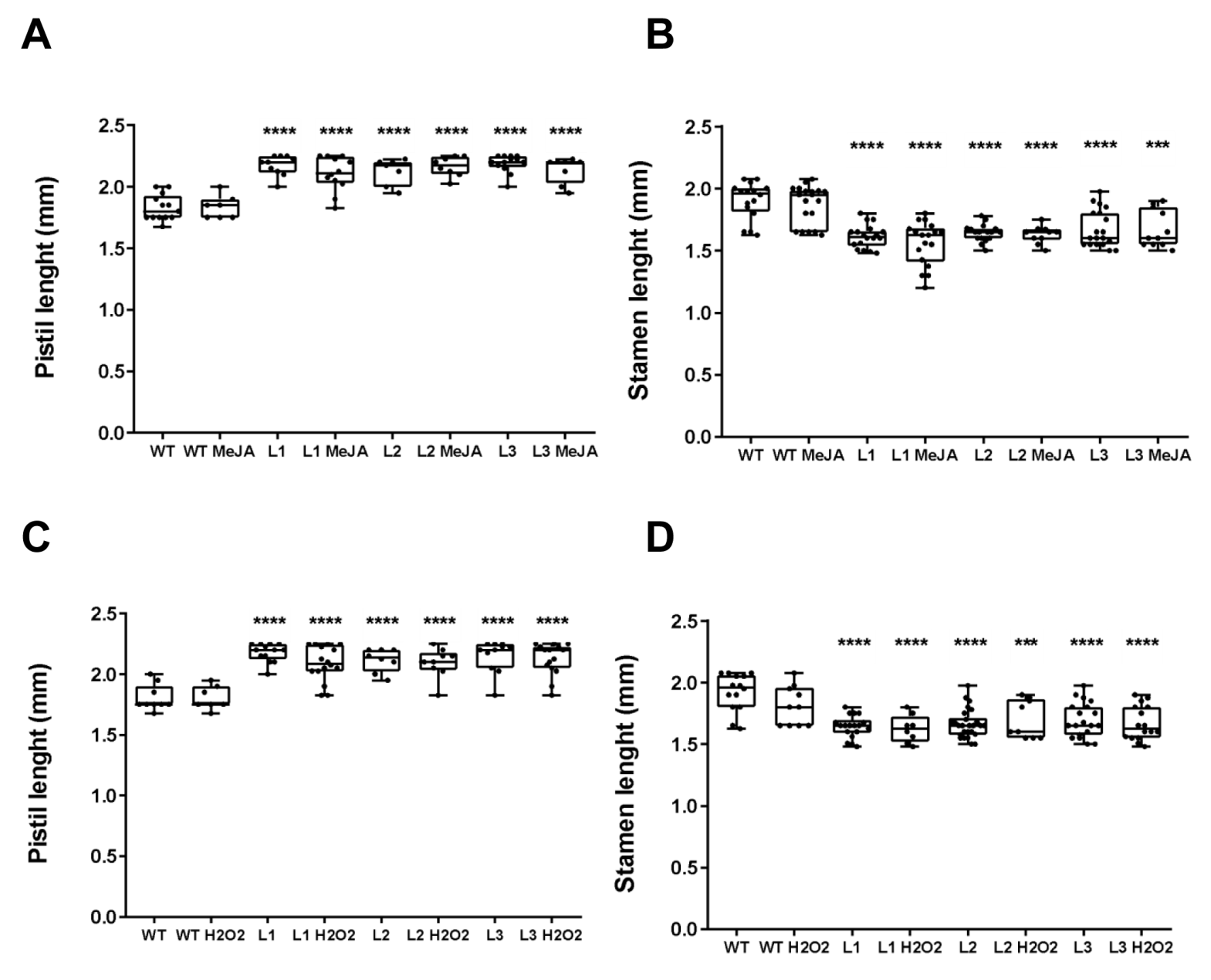

**Fig. S2. Indehiscence of anthers is rescued by Methyl Jasmonate (MeJA) exogenous application in amiR-VLG plants.**

Flowers in 4-week-old WT and amiR-VLG (lines L1, L2 and L3) plants were treated with 1 mM MeJA for three days. Three days after the last application pistil **(A)** and stamen **(B)** lengths were measured (n ≥ 20 flowers from each line).

Flowers in 4-week-old plants WT and amiR-VLG (lines L1, L2 and L3) plants were treated with 1 mM H_2_O_2_ for three days. Three days after the last application pistil **(C)** and stamen **(D)** lengths were measured (n ≥ 20 flowers from each line).

A) Flowers of amiR-VLG plants were treated with 0.5 mM JA for consecutive three days. Arrow shows the moment treatment was applied. B) Silique length of JA treated plants. Asterisks indicate significant difference compared to control plants. ANOVA and Tukey's multiple comparisons test, p < 0.0001 (****). Scale bar: 10 mm.

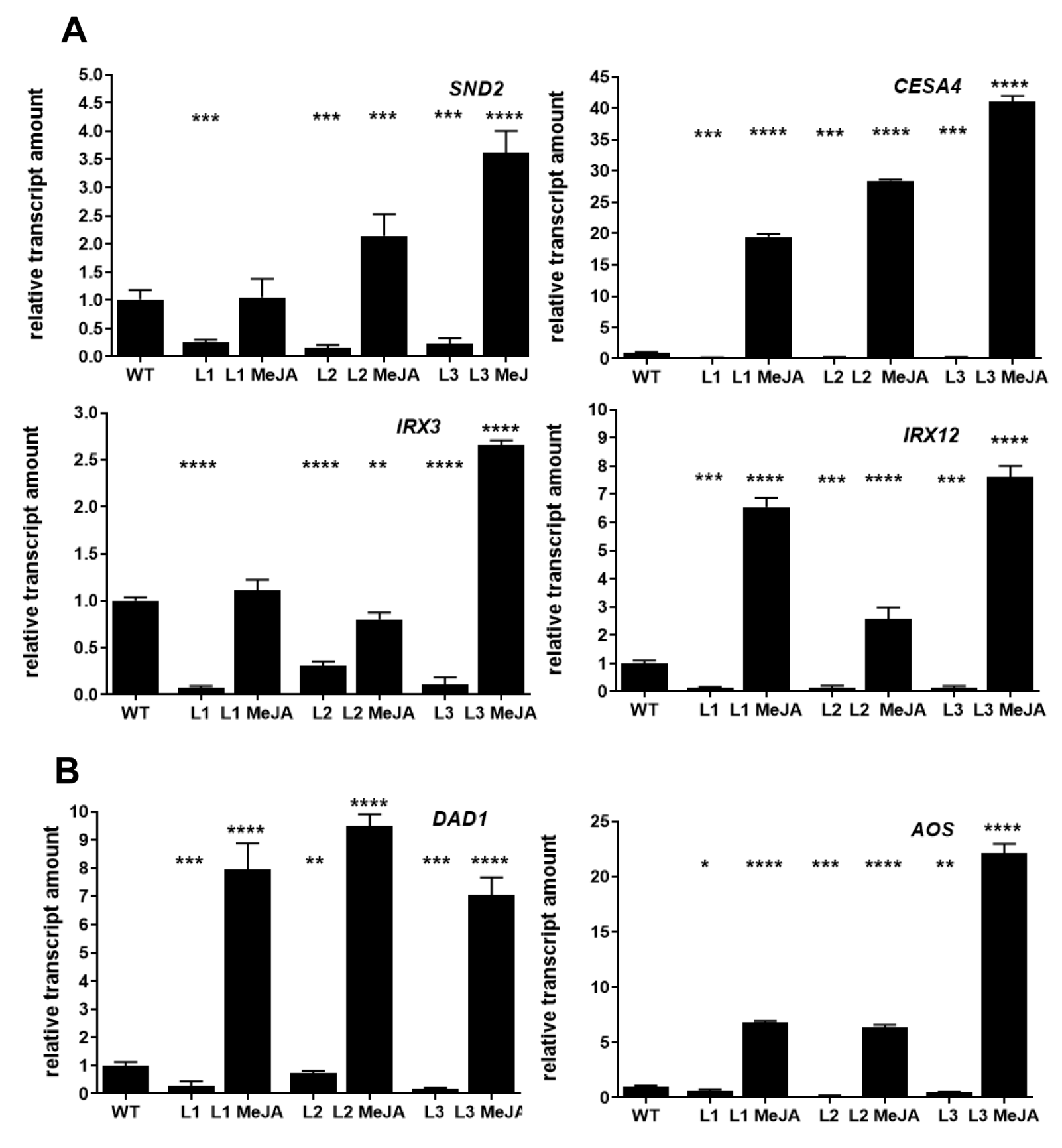

**Fig. S3 Transcript levels from lignin and JA biosynthesis pathways are affected by MeJA exogenous application in amiR-VLG lines.** Relative transcript levels were determined by RT qPCR from total RNA samples isolated from anthers (pooled from stages 1 to 11) of 5-week-old WT and amiR-VLG lines L1, L2 and L3 treated with 1 mM MeJA for three days. Genes from lignin biosynthesis pathway: *SND2 (Secondary wall-associated NAC Domain 2*), *CESA4* (*Cellulose synthase A4*), *IRX3 (Irregular Xylem 3)* and *IRX12 (Irregular Xylem 12)* **(A)**. Genes from JA biosynthesis pathway: *DAD1* (*Defective in Anther Dehiscence 1)* and *AOS* (*Allene* *Oxide Synthase)* **(B)**. Data are means of three replicates ± SD. Asterisks indicate significant differences in comparison to WT plants by ANOVA and Dunnett's multiple comparisons test, p<0.05 (*), p<0.01 (**), p<0.001 (***), p < 0.0001 (****).

Fig. S3

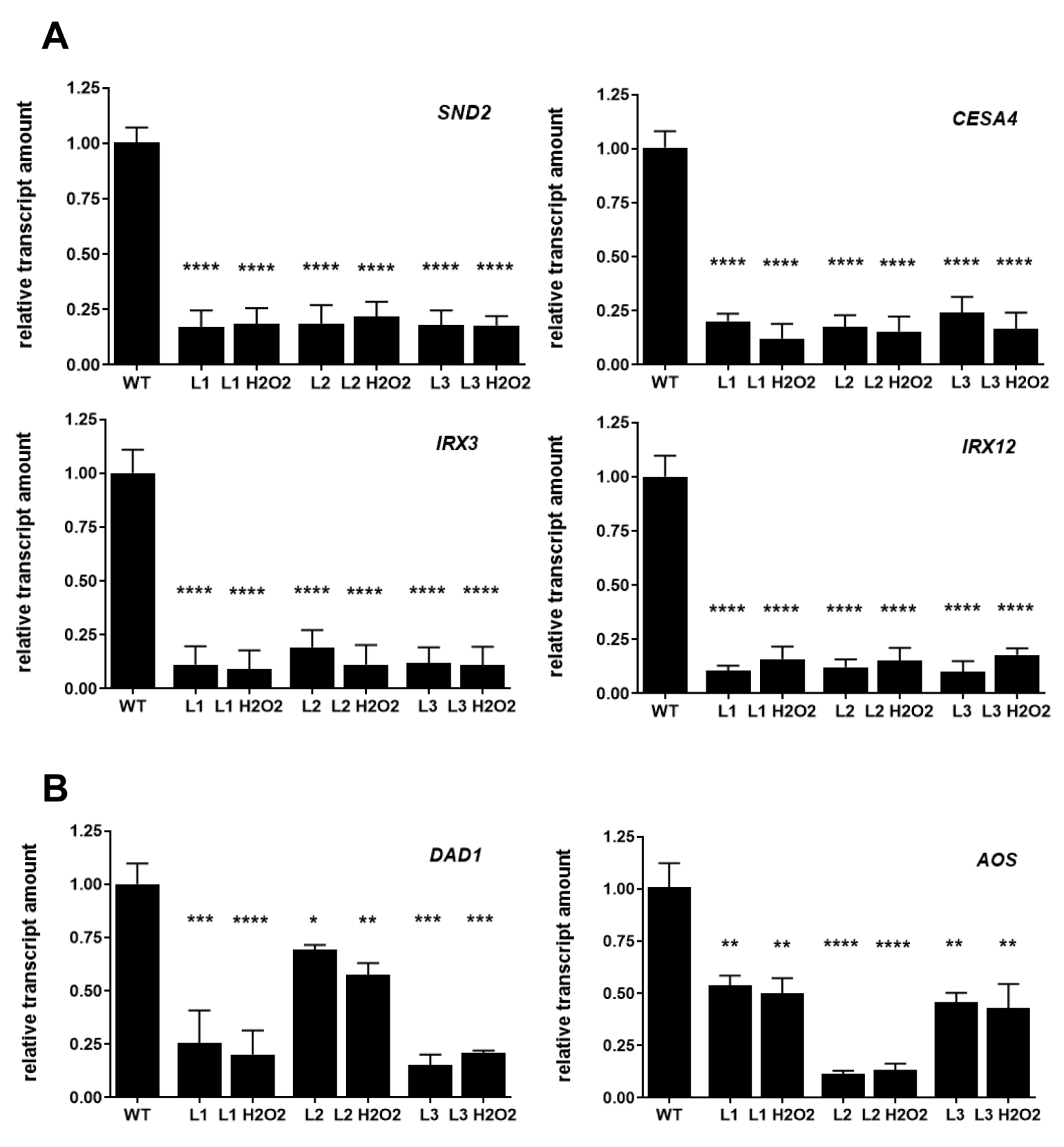

**Fig. S4 Transcript levels from lignin and JA biosynthesis pathways are not affected by H_2_O_2_ exogenous application in amiR-VLG lines.** Relative transcript levels were determined by RT qPCR from total RNA samples isolated from anthers (pooled from stages 1 to 11) of 5-week-old WT and amiR-VLG lines L1, L2 and L3 treated with 1 mM H_2_O_2_ for three days. Genes from lignin biosynthesis pathway: *SND2 (Secondary wall-associated NAC Domain 2*), *CESA4* (*Cellulose synthase A4*), *IRX3 (Irregular Xylem 3)* and *IRX12 (Irregular Xylem 12)* **(A)**. Genes from JA biosynthesis pathway: *DAD1* (*Defective in Anther Dehiscence 1)* and *AOS* (*Allene* *Oxide Synthase)* **(B)**. Data are means of three replicates ± SD. Asterisks indicate significant differences in comparison to WT plants by ANOVA and Dunnett's multiple comparisons test, p<0.05 (*), p<0.01 (**), p<0.001 (***), p < 0.0001 (****).

Fig. S4

**Supplementary Table S1. Primers used for *VLG* specific amiRNA generation.**

| Primer | Sequence |
| --- | --- |
| I miR_DC2-s | GATACTGAAGGGCGGGACGCCTTTCTCTCTTTTGTATTCC |
| II miR_DC2-a | GAAAGGCGTCCCGCCCTTCAGTATCAAAGAGAATCAATGA |
| III miR_DC2*s | GAAAAGCGTCCCGCCGTTCAGTTTCACAGGTCGTGATATG |
| IV miR_DC2*a | GAAACTGAACGGCGGGACGCTTTTCTACATATATATTCCT |
| Primer A | CATTTCATTTGGAGAGAACACG |
| Primer B | CGAAACCGATGATACGAACG |

**Supplementary Table S2. Primers used for q-PCR**

| Primer | Sequence | Reference |
| --- | --- | --- |
| VLG-F | GAAATGTGACTACGGTGCTCA | This work |
| VLG-R | CCACTTCCTCCTTCTCCTCCT |  |
| ACT2-F | GCCATCCAAGCTGTTCTCTC | (Soto et al. 2015) |
| ACT2-R | GAAACCCTCGTAGATTGGCA |  |
| SND2-F | TGATGAAGTTGTGAGCACTGAA | (Hussey et al. 2011) |
| SND2-R | TGCGTCATCTCTTACCTTGC |  |
| CESA4-F | TTGGTGTTGTTGCCGGAGTT | (Hussey et al. 2011) |
| CESA4-R | AACAGTCGACGCCACATTGC |  |
| IRX3-F | GGCAAACTCAAGTGGCTTGAGCG | (Mitsuda et al. 2007) |
| IRX3-R | TAACTCCGCTCCATCTCAATTCC |  |
| IRX12-F | GGTGGATGGGTCGTCATGAGATTC | (Mitsuda et al. 2007) |
| IRX12-R | CGTGGCGTGATGTTGATATGTCGCCC |  |
| AOS-F | TCCGATTTCTCTCCACCCAAA | (Hickman et al. 2017) |
| AOS-R | TGACCCGGAAGCTTTGATCG |  |
| DAD1-F | CTCCTTGAGAAGCAAGGCACGAAG | (Ishiguro et al. 2001) |
| DAD1-R | CCGAAGCTCCTTACCGATTTTCAG |  |
